# The Neural Oscillatory Basis of Perspective-Taking in Autistic and Non-Autistic Adolescents using MEG

**DOI:** 10.1101/2024.10.23.619814

**Authors:** Robert A. Seymour, Gina Rippon, Gerard Gooding-Williams, Hongfang Wang, Klaus Kessler

## Abstract

Taking another’s perspective is a high-level mental skill underlying many aspects of social cognition. Perspective-taking is usually an embodied egocentric process whereby people mentally rotate themselves away from their physical location into the other’s orientation. This is accompanied by increased theta-band (3–7 Hz) brain oscillations within a widespread fronto-parietal cortical network including the temporoparietal junction. Individuals with autism spectrum disorder (ASD) have been reported to experience challenges with high-level perspective-taking, particularly when adopting embodied strategies. To investigate the potential neurophysiological basis of these autism-related individual differences, we used magnetoencephalography in combination with a well-replicated perspective-taking paradigm in a group of 18 autistic and 17 age-matched non-autistic adolescents. Findings revealed that increasing the angle between self and other perspective resulted in prolonged reaction times for the autistic group during perspective-taking. This was accompanied by reduced theta power across a wide network of regions typically active during social cognitive tasks. On the other hand, the autistic group showed greater alpha power decreases in visual cortex compared with the non-autistic group across all perspective-taking conditions. These divergent theta and alpha power effects, coupled with steeper response time slopes, suggest that autistic individuals may rely more on alternative cognitive strategies, such as mental object rotation, rather than an egocentric embodied approach. Finally, no group differences were found when participants were asked to track, rather than take, another’s viewpoint, suggesting that autism-related individual differences are specific to high-level perspective-taking.

## Introduction

Humans possess highly developed social skills that allow us to imagine what others might be experiencing, thinking or feeling to an extent not shared by other species (Tomasello et al., 2005). One important aspect of this is the ability to understand another’s visuospatial experience of the world – a skill termed visuospatial perspective taking, VPT (Flavell et al., 1981). In this study, we used MEG to investigate the neurocognitive basis of perspective-taking between autistic adolescents and age-matched non-autistic participants.

### Perspective-Taking versus Perspective Tracking

Considering another’s visuospatial perspective of the world can take one of two forms or levels (Flavell, Everett, Croft, & Flavell, 1981). First, one can track another’s view (“can the other person see the object?”), sometimes termed level 1 perspective-taking, VPT-1. Second, one can understand *how* the world looks from another’s point of view (“how does the world look like to the other person?”), also termed level 2 perspective-taking, VPT-2. Experimentally, perspective-*tracking* (VPT-1) is often tested by employing visibility judgements (e.g. “is the target visible or occluded from the other’s viewpoint?”, Michelon & Zacks, 2006; Kessler & Rutherford, 2010; for review Samuel et al., 2024), while perspective-*taking* (VPT-2) is typically measured by asking participants how a visual scene would appear from another’s distinct visual perspective (e.g. “is the object to the left or right from the other’s viewpoint?” e.g. Michelon & Zacks, 2006; Hamilton, Brindley & Frith, 2009; Kessler & Rutherford, 2010, for review Samuel et al., 2024). The two processes have been related to different developmental stages, with perspective-tracking emerging around age 2, and perspective-taking emerging around ages 4-5 (Flavell et al., 1981; Gzesh & Surber, 1985; Moll & Tomasello, 2006). Furthermore perspective-tracking has been observed in other species such as apes and corvids (Bugnyar et al. 2004; Bräuer et al. 2007), whereas perspective-taking seems to be uniquely human (Karg et al., 2016).

There is growing evidence that perspective-taking (VPT-2) typically recruits an embodied cognitive process, grounded in the internal bodily and action representations of the observer. Using posture manipulations, several studies (Gooding-Williams et al., 2017; Kessler & Rutherford, 2010; Kessler & Thomson, 2010; Surtees et al., 2013; Wang et al., 2016) have shown that perspective-taking engages large parts of the neuronal bases of the body schema, i.e. the cortical correlates of the internal representation of the body (Coslett et al., 2008; Medina et al., 2009), with some studies also showing that this embodied process is only engaged during VPT-2 but not during VPT-1 (Kessler et al., 2014; Kessler & Rutherford, 2010; Martin et al., 2020; Wang et al., 2016). This suggests that high-level perspective-taking involves the simulated rotation of the embodied self into another’s orientation and perspective (Kessler & Thomson, 2010; Surtees et al., 2013; Wang et al., 2016). In other words, humans literally rotate their own perspective to understand another’s viewpoint. It is important to note that embodied mental self-rotation is not engaged during level-1 perspective-tracking, which seems to recruit a simpler process of inferring another’s line of sight (Kessler et al., 2014; Kessler & Rutherford, 2010; Martin et al., 2020; Zacks & Michelon, 2005); for a recent discussion see Samuel et al., (2024). Interestingly, there are individual differences in the efficiency of this embodied strategy, with East-Asian participants faster than Western participants, potentially reflecting an egocentric bias in Western cultures (Kessler et al., 2014) and with participants with lower social skills scores showing slower perspective taking times (Brunyé et al., 2012) and less embodiment (Kessler & Wang, 2012), potentially reflecting alternative strategies when computing another’s perspective Kessler & Wang, 2012; Samuel, 2024).

### The Neural Correlates of Embodied Perspective-Taking

Since the advent of modern brain imaging there has been steady progress in characterising the neural correlates of embodied perspective-taking. In particular, the posterior division of the temporo-parietal junction (TPJ) (Igelström, Webb and Graziano 2015; Bzdok et al. 2013), seems to play a key role in perspective-taking and more generally in representing other’s mental states (Bögels et al., 2015; Schurz et al., 2013; Van Overwalle, 2011). Using MEG, Wang et al. (2016) investigated embodied simulation during perspective-taking using a well replicated paradigm (Kessler & Rutherford, 2010). In contrast to perspective-*tracking (visibility judgements)*, increases in low-frequency theta-band oscillations (3-7Hz) were found for both the cognitive effort (amount of angular disparity between self vs. other’s viewpoint) and for embodied processing (posture congruence) during perspective-*taking (directionality judgements)*. These effects converged in the right pTPJ for perspective-taking, while for tracking the frontal eye fields were primarily involved, corroborating previous research (Grosbras et al., 2005; Wallentin, 2012; Wallentin et al., 2008) in support of a gaze-based line-of-sight mechanism.

Brain stimulation has also been used to add causal evidence to these neuroimaging findings. Using disruptive dual-pulse TMS, Wang et al., (2016) demonstrated that interfering with the TPJ significantly reduced congruent posture effects, strongly supporting the role of embodiment in perspective-taking. Martin et al., (2020) used HD-tDCS to target the right TPJ during perspective-taking and tracking, reporting an increased posture congruency effect for perspective-taking during stimulation compared to SHAM, but no such effect for VPT-1. In a further TMS study, Gooding-Williams et al. (2017) used repetitive TMS entrainment over right pTPJ to show that TMS pulses administered at theta frequency (6Hz) accelerated perspective-taking, while alpha (10Hz) entrainment slowed perspective-taking down.

Theta rhythms may therefore be the crucial neural frequency to facilitate brain connectivity between the TPJ and other brain regions involved in perspective-taking. In support of this Seymour et al. (2018) reported top-down directional connectivity from prefrontal areas to right TPJ, as well as increased phase-coupling at theta frequencies between right TPJ and the wider social brain network (e.g. ventromedial prefrontal cortex and medial parietal cortex (Alcalá-López et al., 2018; Lieberman, 2007). Wang et al., (2016) also reported increases in theta-band power for the lateral PFC during the cognitive effort of perspective-taking, which was replicated by Seymour et al (2018) and extended to include anterior cingulate cortex (ACC). Activity within these regions during social cognition has been argued to reflect high-level reasoning and working memory processes recruited more generally during complex perspective-taking and mentalizing tasks (for a meta-analysis see Van Overwalle, 2011). Overall there is emerging evidence that perspective-taking is underpinned by an embodied cortical network centred on the right TPJ, as well as including the wider social brain network, body schema and executive control regions, coordinated via theta rhythms (Seymour et al., 2018)

### Perspective-Taking in Autism Spectrum Disorder

In this study we investigated the neural basis of perspective-taking in participants diagnosed with autism spectrum disorder (ASD) – a neurodevelopmental condition characterised by difficulties in social interaction, language, repetitive behaviours and sensory sensitives (APA, 2013). Research in this area has used a mixture of methodologies and produced ambiguous results, with some studies reporting reduced perspective-taking accuracy in autistic children (Baron-Cohen, 1989; Hamilton et al., 2009; Yirmiya, Sigman, & Zacks, 1994), whereas others suggest performance on-par with non-autistic control participants (Hobson, 1984; Tan & Harris, 1991; Leslie & Frith, 1988). Reviewing the literature, Pearson et al., (2013) suggest that overall, ASD participants display a selective impairment for embodied perspective-*taking* (VPT-2), but not perspective *tracking* (VPT-1).

Impaired perspective-taking could also link with so-called “mindblindness” theories of autism (Baron-Cohen, 1997). It has been argued that embodiment scaffolds the development of high-level social cognitive functions like theory of mind (Aichhorn et al., 2006; Hamilton, 2009). Taking another’s viewpoint involves transforming our own sense of self onto another, not in a physical sense as per visual perspective-taking, but in a highly conceptual manner (Hamilton, 2009). Impaired visual perspective-taking could therefore impact the development of social skills in autism more generally (Hamilton et al., 2009; Pearson et al., 2016; Surtees et al., 2013), especially theory of mind. This also raises the intriguing possibility that perspective-taking interventions in ASD may transfer into beneficial outcomes for high-level social and cognitive development (Pearson, 2016).

If autistic individuals fail to use embodiment, could they turn to alternative compensatory strategies? One study suggests so, with autistic participants favouring a mental object rotation strategy during perspective-taking (Pearson et al., 2016). In other words, autistic participants tend to mentally rotate the world towards their own perspective. This approach leads to the correct answer, but is more cognitively demanding than embodiment, resulting in longer reaction times and decreased accuracy (Kessler & Thomson, 2010). In support of this idea, Kessler & Wang (2012) reported that participants with higher levels of autistic traits, but not diagnosed with autism, showed reduced embodiment effects, compared to participants with lower levels of autistic traits. That is, adults with more autistic traits appeared to be less inclined to complete tasks by mentally rotating their body schema into the target orientation. Instead, these individuals were more likely to select an alternative method to embodied transformation such as mentally rotating the target objects (allocentric rotation; Shepard & Metzler, 1988) or making a visuospatial transposition judgement in form of “my left is their right” at high angular disparities (e.g. “spatial transposers”, Gardner et al., 2013; Kessler & Wang, 2012). Accordingly, Brunyé et al., (2012) also showed increased response times for participants with stronger autistic traits. While impaired embodied perspective-taking in autism has been reported behaviourally (Pearson et al., 2013), no study to date has investigated the oscillatory neural correlates of perspective-taking and perspective-tracking in autism.

### Current Study

The current study aimed to investigate autism-related differences in the neurocognitive mechanisms underlying perspective-taking, using MEG. We collected data from a group of 18 autistic adolescent participants and 17 age-matched controls, using the same perspective-taking paradigm and data analysis pipeline as in Seymour et al. (2018). It was hypothesised that the adolescent control group would adopt an embodied egocentric strategy during perspective-taking, resulting in increased theta-band power (3-7Hz) within the TPJ, prefrontal executive control regions and regions coding for body schema, as per Wang et al., (2016) and Seymour et al. (2018). However, for the autistic group it was hypothesised that alternative strategies would be preferred. That is, either an object mental rotation strategy (Conson et al., 2015; Pearson et al., 2016; Kessler & Wang, 2012) or a visuospatial transposition strategy (“my left is their right”) would be adopted (Gardner et al., 2013; Kessler & Wang, 2012). Especially an object rotation strategy would result in longer reaction times (e.g. Kessler & Thomson, 2010) and oscillatory signatures in the alpha frequency range reflecting increased processing in visual/parietal cortex (Gardony et al., 2017; Hanslmayr et al., 2005; Kasten & Herrmann, 2017; Klimesch et al., 2003; Riečanský & Katina, 2010). This hypothesis was additionally motivated by neurostimulation research showing enhanced mental object rotation performance during alpha-band repetitive TMS (Klimesch et al., 2003), as well as alpha-specific TACS stimulation (Kasten & Herrmann, 2017). A visuospatial transposition strategy should also result in stronger alpha signatures, given alpha’s pivotal role in visuospatial attention (Sauseng et al., 2005). Finally, it has suggested that perspective-taking difficulties in autism are specific to VPT-2 (Pearson et al., 2013, 2016), thus, we hypothesised no significant oscillatory differences in perspective-tracking (VPT-1) between autistic and non-autistic groups.

## Materials and Methods

### Participants

Data were collected from 18 autistic participants and 17 age-matched typically developing controls, see Table 1. MEG data from 4 of these participants (2 ASD, 2 control) were excluded based on RT results (see *Behavioural Data Analysis*). Autistic participants had a confirmed clinical diagnosis of ASD or Asperger’s syndrome from a paediatric psychiatrist. Participants were excluded if they were taking psychiatric medication or reported epileptic symptoms. Non-autistic participants were excluded if a sibling or parent was diagnosed with autism.

**Table 1.** Participant demographic and behavioural data. * = behavioural scores significantly greater in Autistic > Non-Autistic group, t-test, p<.05.

|  | <u>N</u> | <u>Age</u> | <u>Male/Female</u> | <u>Autism</u><br><u>Quotient</u><br><u>(Adult)</u> | <u>Raven</u><br><u>Matrices</u><br><u>Score</u> | <u>Glasgow</u><br><u>Sensory</u><br><u>Score</u> | <u>Mind</u><br><u>in the</u><br><u>Eyes</u><br><u>Score</u> |
| --- | --- | --- | --- | --- | --- | --- | --- |
| **Autistic group** | 18 | 16.67 | 14 male; 4 female | 32.6* | 43.8 | 65.3* | 21.8 |
| **Comparison group** | 17 | 16.41 | 14 male; 3 female | 10.71 | 48.8 | 38.4 | 25.5 |

All experimental procedures complied with the Declaration of Helsinki and were approved by the Aston University, Department of Life & Health Sciences ethics committee. Written consent was obtained from participants aged 18 or over, or a parent/guardian for participants aged under 18.

### Experimental Paradigm and Design

The paradigm was adapted from Kessler and Rutherford (2010). The stimuli were coloured photographs (resolution of 1024 × 768 pixels), showing an avatar seated at a round table from one of four possible angular disparities (see Figure 1A). In each trial, one of the grey spheres on the table turned red indicating this sphere as the target. From the avatar’s viewpoint, the target could be either visible/occluded (VO) by a centrally presented black screen (perspective-tracking); or to the left/right (LR) (perspective-taking). Stimuli were presented in 12 mini-blocks of 32 trials, alternating between LR and VO conditions. On each trial, participants were asked to make a target location judgement according to the avatar’s perspective by pressing the instructed key on an MEG-compatible response pad: the left key for “left” or “visible” targets from the avatar’s viewpoint and the right key for “right” or “occluded” targets. Accuracy feedback was provided after each trial in the form of a short tone. As in Kessler and Rutherford (2010), we collapsed across clockwise and anticlockwise disparities, and separately collapsed correct responses for left and right and visible and occluded, respectively. This resulted in four separate experimental conditions (for examples see Figure 1): left/right judgements where the avatar is 160^0^ from own perspective (LR-160); left-/right judgements where the avatar is 600 from own perspective (LR-60); visible/occluded judgments where the avatar is 1600 from own perspective (VO-160); visible/occluded judgments where the avatar is 600 from own perspective (VO-60). This 2×2 design allowed us to disentangle perspective-taking from perspective-tracking and investigate the effect of an increased angular disparity, which has been shown to lengthen reaction times for perspective-taking but not for perspective-tracking (Kessler & Rutherford, 2010; Michelon & Zacks, 2006). We chose to use 160° vs. 60° based on the results of Seymour et al. (2018), Gooding-Williams et al., (2017) and Wang et al., (2016).

**Figure 1.**
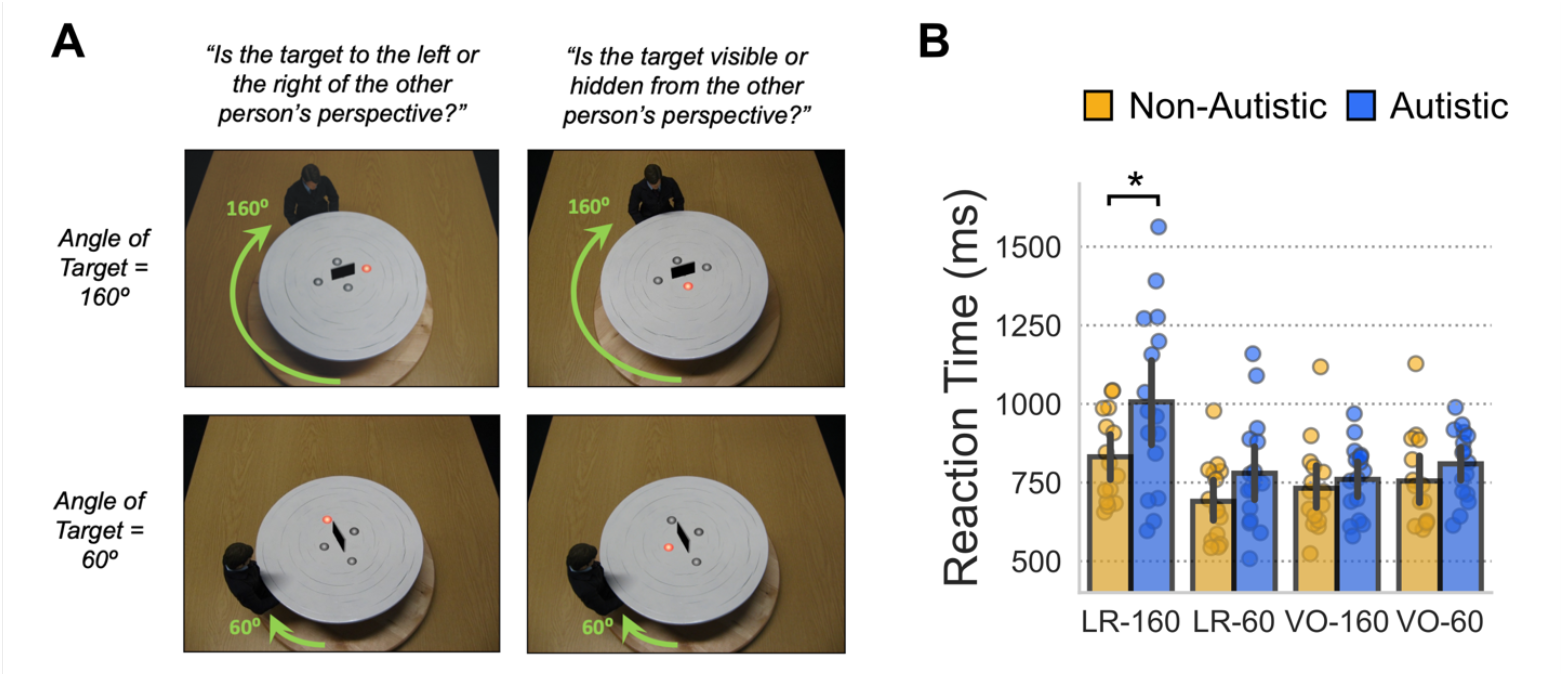
**(A)** Example stimuli from the perspective taking paradigm, for each of the four experimental conditions. Arrows are for illustrative purposes and weren’t presented to participants. **(B)** Reaction time (RT) data are shown for each of the four experimental conditions. Individual datapoints indicate participant RT medians per condition. Significant differences, p<.05, between groups were found for the LR-160 condition.

### Behavioural Data Analysis

All trials containing incorrect answers or response times greater than 2 standard deviations from the participant’s individual median reaction times (across all experimental conditions) were excluded from subsequent analyses. Median RT per condition from each participant, as per Kessler & Rutherford (2010), were entered into a 2×2×2 mixed ANOVA with condition (LR, VO) and angle (160°, 60°) as repeated measures factors and group (ASD, control) as a between-subjects factor, using the JASP statistics package. Data from 4 participants (2 ASD, 2 control) were discarded due to a median RT greater than 2 standard deviations from the group median.

### MEG and Structural MRI Acquisition

MEG data were acquired using a 306-channel Neuromag MEG scanner (Vectorview, Elekta, Finland) made up of 102 triplets of two orthogonal planar gradiometers and one magnetometer. All recordings were performed inside a magnetically shielded room at a sampling rate of 1000Hz. Five head position indicator (HPI) coils were applied for continuous head position tracking, and visualised post-acquisition using an in-house Matlab script. For MEG-MRI coregistration purposes three fiducial points, the locations of the HPI coils and 300-500 points from the head surface were acquired using the integrated Polhemus Fastrak digitizer. Visual stimuli were presented on a projection screen located 86cm from participants, and auditory feedback through MEG-compatible headphones. Data acquisition was broken down into three sequential runs, each lasting 8-10 minutes.

A structural T1 brain scan was acquired for source reconstruction using a Siemens MAGNETOM Trio 3T scanner with a 32-channel head coil (TE=2.18ms, TR=2300ms, TI=1100ms, flip angle=9°, 192 or 208 slices depending on head size, voxel-size = 0.8×0.8×0.8cm).

### MEG Preprocessing

All MEG data were pre-processed using Maxfilter (temporal signal space separation, .9 correlation), which supresses external sources of noise from outside the head (Taulu and Simola 2006). To compensate for head movement between runs, data from runs 2 and 3 were transformed to participant’s head position at the start of the first block using the *-trans* option of Maxfilter. For each participant, the entire recording was band-pass filtered between 0.5-250Hz (Butterworth filter) and band-stop filtered to remove residual 50Hz power-line contamination and its harmonics. Data were then epoched into segments of 2.5s (1s pre, 1.5s post stimulus onset) and each trial was demeaned and detrended. Trials containing artefacts (SQUID jumps, eye-blinks, head movement) were removed by visual inspection, resulting in removal of an average of 86.5 trials per condition, per participant. Four MEG channels containing large amounts of non-physiological noise were removed from all source-level analyses. The pre-processed data were then separated into experimental conditions and downsampled to 200Hz to aid computation time.

### MEG-MRI Coregistration

MEG data were co-registered with participants’ T1 MRI structural scan by matching the digitised head shape data with surface data from the structural scan (Jenkinson & Smith, 2001). Subsequently, the aligned MRI-MEG image was used to create (1) a forward model based on a single-shell description of the inner surface of the skull (Nolte, 2003), using the segmentation function in SPM8 and (2) spatial normalisation parameters to create individual volumetric grids. To facilitate group analysis, each individual volumetric grid was warped to a template based on the MNI brain, 8mm resolution. Subsequently the inverse of the normalisation parameters were applied to the template grid, for source analysis.

### Sensor Level Analysis

Sensor-level time-frequency representations (TFRs) were calculated using a single Hanning taper between frequencies of 2-30Hz in steps of 1Hz. The entire 2.5s epoch was used, with a sliding window of 0.5s, but the first 0.25s and last 0.5s of each trial were discarded to avoid edge artefacts. Due to different scales between the two MEG sensor-types, only data from the gradiometers were used, with TFR power averaged across each adjoining pair of gradiometers post-hoc. All analyses were computed on single trials and subsequently averaged, and therefore TFRs contain both phase-locked (evoked) and non-phase-locked (induced) information.

We focussed our analysis on two frequency bands - theta-band (3-7Hz) and alpha (8-12Hz) power as these have been shown to underly perspective-taking and perspective-tracking on the same task (Seymour et al., 2018; Wang et al., 2016). As per our previous analyses (Seymour et al., 2018) we compared TFRs in relation to angular disparity effects (160^0^ vs 60^0^) in LR and VO trials separately. Next, due to our hypothesis regarding differing cognitive strategies underlying perspective-taking in autistic versus non-autistic participants, we examined TFR differences between groups separately for LR-160 and LR-60 trials. Group-differences related to these contrasts were calculated using cluster-based permutation testing (see *Statistical Analysis*).

### MEG Source-Level

Source localisation was conducted using Linearly Constrained Minimum Variance (LCMV) beamformer (Van Veen et al., 1997) which applies a spatial filter to the MEG data at every voxel of a canonical 0.8 cm brain-grid, in order to maximise signal from that location and attenuating signals elsewhere. The spatial filter was calculated using a covariance matrix from a time–frequency tile centred on the effects found at sensor level. For all analyses, a common filter across baseline and task periods (−0.65s to 0.65s) was used. Due to rank reduction following Maxfilter, a regularisation parameter of lambda 5% was applied to the covariance matrix.

For statistical testing, cluster-based non-parametric permutation testing was used to correct for multiple comparisons across voxels (Maris & Oostenveld, 2007). The resulting whole-brain statistical maps were presented on a cortical mesh using the Connectome Workbench software (Van Essen et al., 2012).

### Statistical Analysis

For MEG data, statistical analysis was performed using cluster-based permutation tests (Maris & Oostenveld, 2007), which consist of two parts: first an independent-samples t-test is performed, and values exceeding an uncorrected 5% significance threshold are grouped into clusters. The maximum t-value within each cluster is carried forward. Second, a null distribution is obtained by randomising the condition label (e.g. Autistic/Non-Autistic) 5000 times and calculating the largest cluster-level t-value for each permutation. The maximum t-value within each original cluster is then compared against this null distribution, and the null hypothesis is rejected if the test statistic exceeds a threshold of p<.05.

### Correlations with Other Measures

We correlated group-differences in the oscillatory basis of perspective-taking with various behavioural data from the autistic group: Autism Quotient (Baron-Cohen et al., 2001); the Glasgow Sensory Questionnaire (Robertson & Simmons, 2013); and reaction time from the perspective-taking behavioural data (LR-160 condition). The Pearson correlation coefficient was used.

The participants in this study also completed a separate MEG scanning session where they viewed visual gratings. These data were previously reported in Seymour et al., (2019) – we found reduced phase-amplitude coupling (PAC) and reduced feedback connectivity in autistic participants. To determine links between these two datasets we correlated group-differences in the oscillatory basis of perspective-taking with PAC from the visual grating dataset as well as 8-12 Hz directed asymmetry index (DAI) which is a measure of feedback vs. feedforward connectivity in the visual system (see Seymour et al., 2019 for more details).

## Results

### Behavioural Results

Median reaction times (RT) from each participant, see Figure 1B, were entered into an 2×2×2 mixed ANOVA, with condition (LR, VO) and angle (160°, 60°) as repeated measures factors and group (ASD, control) as a between-subjects factor. Results showed a significant interaction between angle and condition on RT, F(1,29) = 29.12, p<.001, see Figure 1B, replicating previous studies (Kessler & Rutherford, 2010; Michelon & Zacks, 2006; Seymour et al., 2018; Wang et al., 2016) that reported a significant angular disparity effect for LR judgements (VPT-2) but not VO judgements (VPT-1). Post-hoc tests revealed that the interaction was indeed due to significantly longer RT for the LR-160 conditions compared with all other conditions (ptukey <.001). There was also a main effect of group on RT, F(1,29) = 5.68, p = .024. While there was no statistically significant three-way interaction (p=.18), a post-hoc analysis showed that the autistic group had longer RT compared with the non-autistic group, specifically in the LR-160 condition (p=.005, all other conditions were p>.05, see Figure 1B).

### Theta-Band (3-7 Hz) Results

We first focussed on sensor-level theta-band power given previous research showing that embodied perspective-taking engages a network of brain regions coordinated by theta-band oscillations (Seymour et al., 2018; Wang et al., 2016). Using the same sensor-level time-frequency pipeline as in Seymour et al., (2018) we compared LR-160 vs. LR-60 trials to investigate angular disparity during perspective-taking. As predicted, in the non-autistic control group there were large increases in theta power (3-7 Hz, 0-0.65 s) in LR-160 versus LR-60 trials. However, this was not the case for the ASD group. Statistically comparing theta power between groups revealed one significant cluster of greater (3-7Hz) theta-band power in the non-autistic group at 0.3-0.65 s in the LR-160 versus LR-60 condition, p<.05 (see Figure 2 top panel).

**Figure 2.**
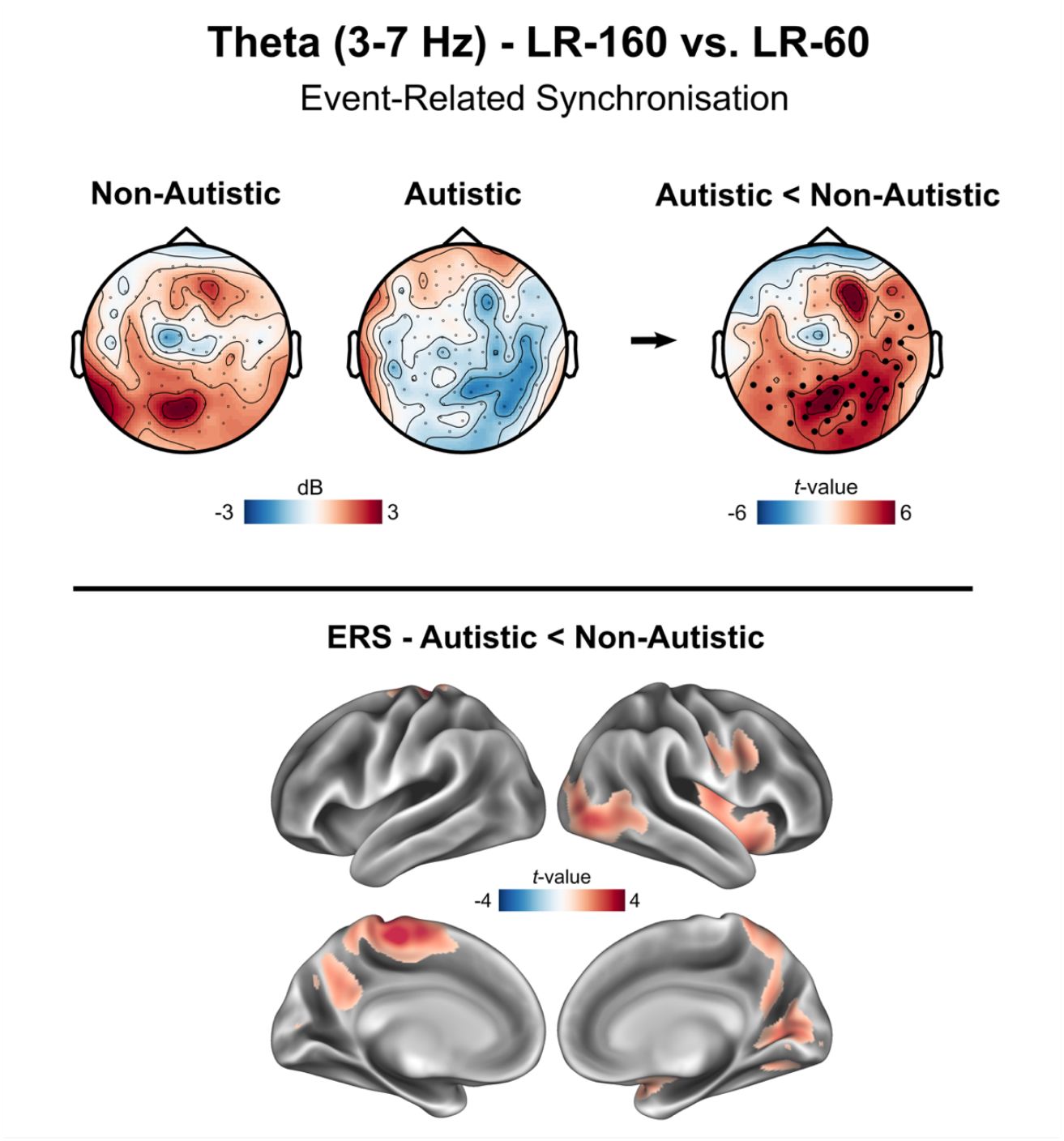
Theta-band (3-7 Hz) perspective-taking results. In the top panel, sensor-level topoplots (gradiometers only) are shown for the difference in theta-band (3-7 Hz, 0-0.65 s) oscillatory power between LR-160 vs. LR-60 trials. Scales represent MEG field strength, with units of decibels (dB). Statistical comparison of the two groups is shown on the far right, with units of t-values. Sensors showing a significant difference between groups, p<.05, are highlighted with a boldened black dot. In the bottom panel the theta-band group difference source-localisation results are shown on an inflated cortical mesh, after correcting for multiple comparisons.

Next, source localisation was performed to find the brain regions underlying these effects. When contrasting LR-160 versus LR-60 trials in the non-autistic group, theta power localised to a widespread collection of regions including ventral visual and occipito-temporal cortex, right temporo-parietal and medial parietal cortex, sensorimotor cortices, as well as lateral, medial, and ventral prefrontal cortex (see Supplementary Figure 1). Statistically comparing 3-7 Hz power in source-space revealed that there were group differences in theta power within left sensorimotor cortex, left precuneus and right lateral prefrontal cortex (Figure 2 – bottom panel).

In contrast to perspective-taking (LR trials), perspective-tracking (VO trials) has previously been shown to rely significantly less on theta-oscillations (Wang et al., 2016). In line with this, when comparing theta power between VO-160 vs. VO-60 trials, both groups showed very small increases in theta-power and no significant group differences were found (see Supplementary Figure 2).

### Alpha-Band (8-12 Hz) Results

Next, we focused on task-related changes in the alpha-band (8-12 Hz) for a number of reasons. First, perspective-taking requires active visual processing which is known to involve modulations of alpha-band power in the occipital lobe (Jensen & Mazaheri, 2010; Klimesch, 1999). Indeed, presentation of the stimuli used in this study are known to robustly elicit alpha-band desynchronization (Seymour et al., 2018; Wang et al., 2016). Second, Wang et al., (2016) found modulation of alpha rhythms during perspective-taking when investigating the effect of angular disparity. Third, given that the autistic group are more likely to adopt a mental rotation rather than embodied strategy (Kessler et al., 2014; Pearson et al., 2016), we hypothesised that the increased visual processing demands would result in greater alpha-related desynchronization versus the control group (Gardony et al., 2017; Klimesch et al., 2003; Riečanský & Katina, 2010).

Sensor-level TFRs were calculated using the same pipeline as in Seymour et al., (2018). For both the ASD and control groups perspective-taking was accompanied by large alpha-band desynchronization over posterior sensors (see Figure 3) in both the LR-160 and LR-60 trials. Statistically comparing groups, we found that the autistic group had greater alpha-desynchronization over posterior sensors in both LR-160 and LR-60 trials (8-12 Hz, 0.2-0.85 s, p < .05).

**Figure 3.**
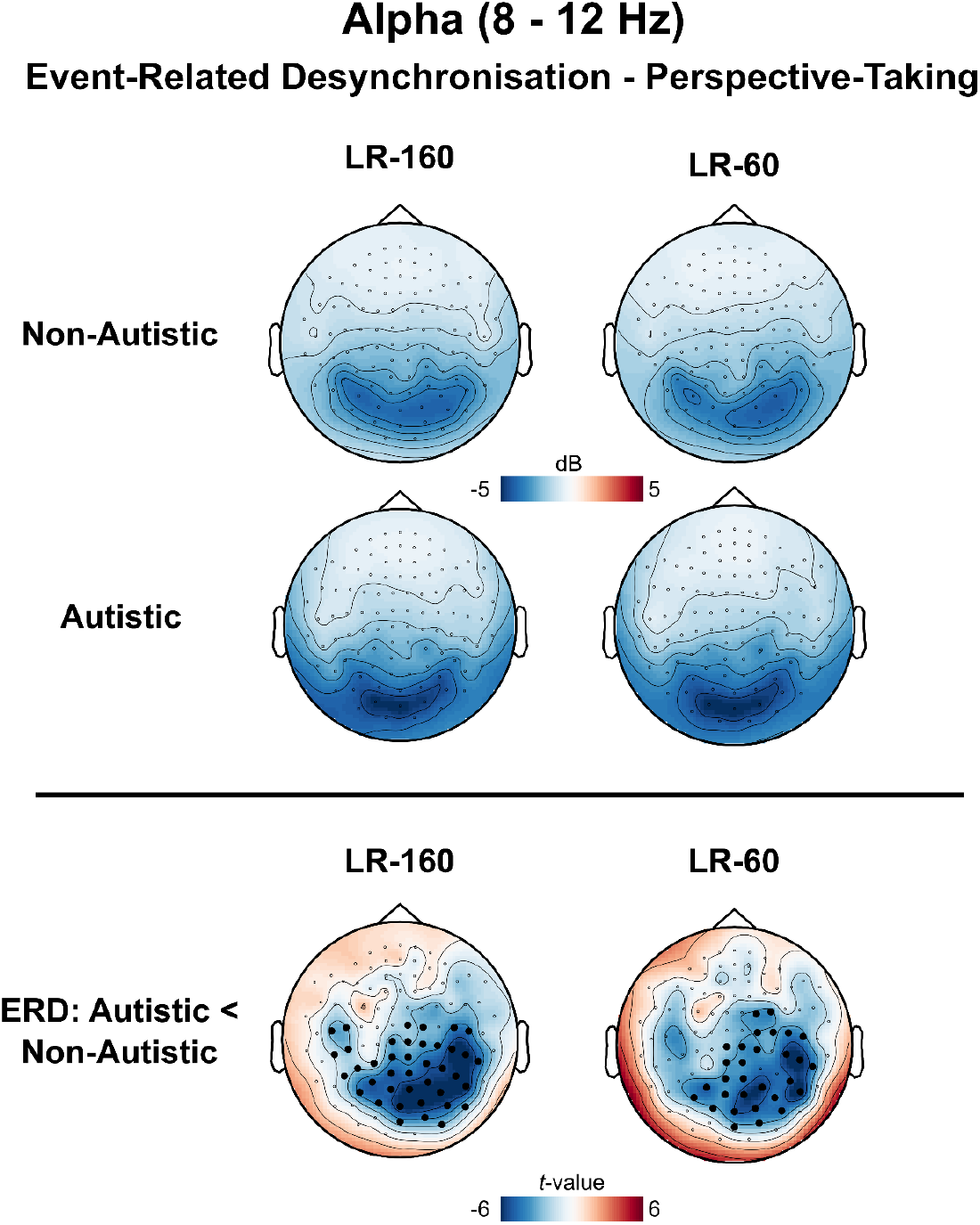
Sensor-level alpha-band (8-12 Hz) perspective-taking results. In the top panel, sensor-level topoplots (gradiometers only) are shown for LR-160 and LR-60 trials (8-12 Hz, 0-0.8 s). Scale represents MEG field strength, baseline-corrected, with units of decibels (dB). Statistical comparison of the two groups is shown in the bottom panel for LR-160 and LR-60 trials separately, with units of t-values, with sensors showing a significant difference, p<.05, highlighted with a boldened black dot.

Interestingly, unlike Wang et al., (2016) we did not observe a statistical difference in alpha-band power when comparing LR-160 versus LR-60 trials in either group and there were also no group x condition interaction effects. The above analyses were also repeated for perspective-tracking (i.e. VO trials), however no group differences, p>.05, were found for alpha-band power.

Using the same source-localisation pipeline as for theta, we investigated the cortical sources underlying 8-12 Hz alpha desynchronisation in perspective-taking. As expected the group difference in alpha power localised to occipital cortex, with a peak in primary visual cortex, in both LR-160 and LR-60 trials, see Figure 4 top panel. To investigate the temporal progression of alpha-related group differences in further detail we extracted virtual time-series from a region of interest in primary visual cortex. In both LR-160 and LR-60 trials the autistic group (yellow line) displayed larger alpha desychronisation from ∼0.35s to 1.0s post-stimulus presentation compared with the non-autistic group (blue line), see Figure 4 bottom panel.

**Figure 4.**
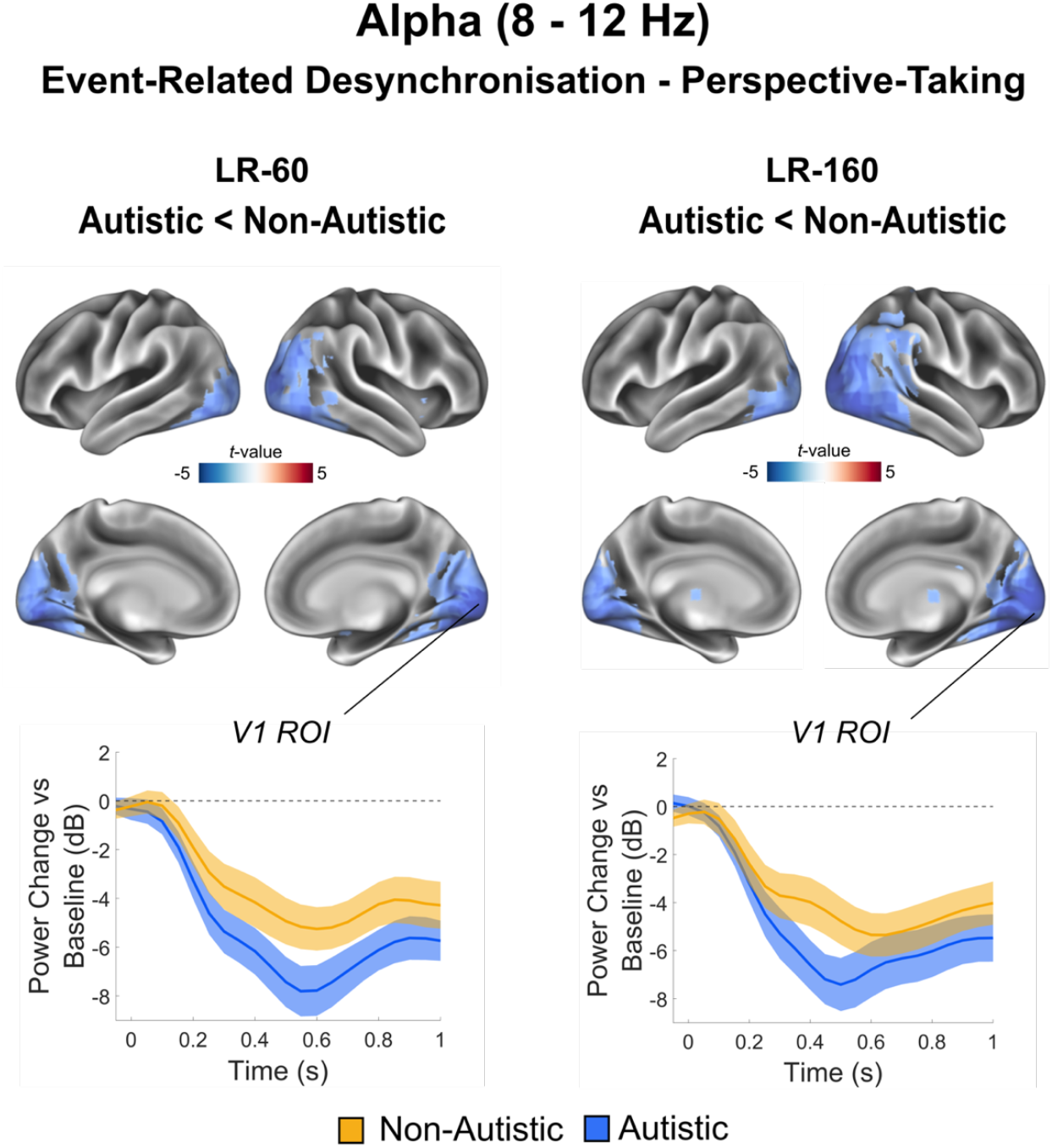
Source-level alpha-band (8-12 Hz) perspective-taking results. Alpha-band power (8-12 Hz, 0-0.8 s) was localised using a beamformer. In the top panel baseline corrected alpha power results are plotted for each condition (LR-160, LR-60) and group (autistic, non-autistic). Only clusters passing a p<.05 corrected threshold are shown. A region of interest was defined in primary visual cortex (V1) and the alpha power change versus baseline was calculated for LR-160 and LR-60 trials. The bottom panel plots this in a time-resolved manner separately for non-autistic (yellow) and autistic groups (blue). Error bars correspond to 95% confidence intervals.

## Discussion

In this study we investigated the neural oscillatory basis of high-level perspective-taking in a group of autistic adolescents and a non-autistic group of age-matched control participants. Behaviourally, the autistic group had longer reaction times compared with age-matched controls, specifically for perspective-taking trials in which the target’s perspective was 160° versus 60° away from the participant’s own perspective (Figure 1B). No group differences were found for perspective-tracking trials. This supports the notion of a selective difference regarding embodied strategies during perspective-taking but not tracking in ASD (Pearson et al., 2013; 2016).

Using MEG, we further showed that when increasing the angular disparity between self and other perspectives (160° versus 60°), non-autistic adolescent participants showed increased theta-band (3-7Hz) power (see Figure 2, top panel). This theta pattern was absent in the autistic group and a direct comparison in source space revealed contrasting theta power differences within ventral visual and occipito-temporal cortex, medial parietal cortex, sensorimotor cortices, and lateral and ventral prefrontal cortex (see Figure 2 bottom panel). However, when examining alpha rhythms a very different picture emerged. Here, there were no group differences when the angular disparity increased from 60° to 160°. Instead across both LR-160 and LR-60 trials the autistic group showed greater alpha desynchronisation over occipital cortices from 0.3s after stimulus onset compared with the non-autistic group.

When examining perspective-tracking (VPT-1) trials, for both, theta and alpha-bands no significant differences between groups were observed, corroborating the null result in behavioural data and the view that autistic and non-autistic individuals do not appear to differ in their strategy for perspective-tracking (Pearson et al., 2013).

The differences in oscillatory signatures observed between the two groups for perspective-taking (VPT-2), i.e. theta versus alpha effects, may reflect differences in cognitive strategies employed during perspective taking (Gardener et al., 2013; Kessler & Wang, 2012; Pearson et al., 2013, 2016; Samuel et al., 2023). Stronger theta power in the non-autistic group localised to ventral visual and occipito-temporal cortex, right temporo-parietal and medial parietal cortex, sensorimotor cortices, as well as lateral, medial, and ventral prefrontal cortex (see Supplementary Figure 2). This replicates previous MEG studies (Seymour et al., 2018; Wang et al., 2016) showing increased theta power in a similar set of brain regions during perspective-taking, coordinated by the right TPJ. We believe this pattern of brain network activation in the non-autistic control group, especially theta activity within right TPJ and sensorimotor cortex, reflects the use of an embodied strategy (Kessler & Thomson, 2010; Wang et al., 2016; Gooding-Williams et al., 2017). The involvement of medial and lateral prefrontal regions is thought to manage conflict between self and other perspectives, which has been shown to be affected in autism (Carrington & Bailey, 2009; Conson et al., 2015; Hartwright et al., 2012). Our findings imply that autistic adolescents are less able to, or simply less inclined to use such an embodied mental self-rotation strategy.

The perspective-taking paradigm used in this study can, however, be solved in another way – through mental object rotation (Kessler & Thomson, 2010; Samuel et al., 2023; Zacks & Tversky, 2005). Alternatively, autistic participants might favour a visuospatial strategy where the other’s left and right are still aligned with the egocentric left and right at low angular disparity (60°), allowing for a very quick left/right decision, while at high angular disparity (160°, i.e. almost face-to face) their egocentric left would be the other’s right and their right the other’s left, only requiring a visuospatial transposition (e.g. Garnder et al., 2013). Our finding of stronger posterior alpha desynchronization hints at the possibility that the autistic group were using one of these alternative strategies to solve the task (Conson et al., 2015; Pearson et al., 2016). Posterior alpha power desynchronisation reflects reduced functional inhibition during visual perception (Jensen & Mazaheri, 2010), suggesting a far more sensory, bottom-up manipulation of the visual stimuli in the ASD group. Furthermore we found a significant negative correlation between alpha desynchronisation and the amount of V4-to-V1 feedback connectivity as measured during a complimentary low-level visual task in the same participants (Seymour et al., 2019) (Supplementary Figure 3). These findings imply that when ASD participants are able to use an alpha-based visual or object-rotation strategy they do so, but at the detriment of the more efficient as well as more socially engaged strategy of embodying another’s perspective through mental self-rotation into their viewpoint. In support of an object-rotation strategy, previous research using TMS (Klimesch et al., 2003), TACS (Kasten & Herrmann, 2017) and neurofeedback (Hanslmayr et al., 2005) have shown that alpha-band oscillations are causally linked with mental object rotation abilities. Interestingly, our alpha desynchronisation group difference did not interact with angular disparity – no effect of alpha power was found when comparing trials where the object was 160° versus 60° from the participant’s perspective. This implies that the same amount of visual processing was engaged by the autistic group despite the 100° difference in rotation required. This could suggest that a visuospatial transposition strategy explanation might be likely (“my left is their right” at 160°) or that the stimuli in this study required only very simple mental rotations along just one dimension rather than true three-dimensional object rotation as often used with Shepard & Metzler type stimuli (Shepard & Metzler, 1988).

More generally, our findings underscore the importance of studying different cognitive strategies in perspective-taking (Kessler et al., 2014; Samuel et al., 2023, 2024). Alpha desynchronisation is likely to be a neural signature of effortful and slower object-rotation strategy or indicative of a visuospatial transposition strategy (e.g. “spatial transposers” Gardner et al., 2013; also Kessler & Wang, 2012), whereas increases in theta power associated with the faster and more embodied self-rotation strategy in controls (Wang et al., 2016). Future research could investigate this further by characterising the neural dynamics of embodied processing and mental rotation within the same participants. Repetitive TMS entrainment could also be used to investigate the causal role of alpha vs theta oscillations as a function of neurocognitive strategy during perspective taking (Gooding-Williams et al., 2017).

Finally, our results show that when instructed to imagine the world from another’s perspective, autistic participants activate visual regions rather than the extended network of regions supporting embodied transformation, including temporal, parietal, sensorimotor, and prefrontal areas (Wang et al., 2016; Seymour et al., 2018). A bias towards prioritising bottom-up sensory information, at the expense of top-down prior information, has been described by predictive-coding accounts of autism (Kessler et al., 2016; Palmer et al., 2017; Pellicano & Burr, 2012). These generally focus on sensory aspects of ASD but can be easily extended for social and embodied processes (e.g. Kessler et al., 2016). Just as one can represent an object for perceptual inference, one can also represent another’s perspective, or mental state, if it helps to predict the causes of sensory input and minimise prediction error (Palmer et al., 2015). Embodied processing may be particularly affected in autism as another’s perspective is abstractly inferred, presumably involving top-down mechanisms and regions at the top of the cortical hierarchy (Friston, 2011). Our findings hint at the intriguing possibility of a common mechanism underlying both sensory and social symptoms in ASD. Impaired embodied abilities may result in cascading deficits or atypicality in the development of social cognitive skills like Theory of Mind (Kessler et al., 2016; Pearson et al., 2013). While the paradigm in this study required a relatively simple spatial judgement (left/right), there is evidence to suggest that visual perspective-taking performance can predict theory of mind in children (Hamilton et al., 2009). For perspective-taking, ASD participants can adopt alternative compensatory strategies, potentially supported by increased alpha desynchronisation over visual cortex (Hanslmayr et al., 2005; Kasten & Herrmann, 2017; Klimesch et al., 2003; Pearson et al., 2016). However, as tasks become more abstract, for example false belief tasks requiring representing what others are thinking (Yuk et al., 2020), this strategy becomes inefficient, if not impossible.

### Limitations

We note three limitations. First, our assertion that alpha desynchronisation reflects mental rotation strategy relies on reverse inference. However, there are both theoretical (Pearson et al., 2013, 2016; Samuel et al., 2023, 2024) and quantitative grounds (Gardony et al., 2017; Hanslmayr et al., 2005; Kasten & Herrmann, 2017; Zacks & Michelon, 2005) to support this. To further our argument, future MEG studies could explicitly ask participants about which strategies they are using, and/or measure mental rotation alongside perspective-taking in the same participants. Second, we did not collect a formal clinical assessment of autism, e.g. the Autism Diagnostic Observation Schedule (Lord et al., 2000), to ascertain symptom severity. Instead, strict participant exclusion criteria were implemented, and only autistic participants with a confirmed clinical diagnosis of ASD or Asperger’s syndrome were included. Between groups, there were significant differences in autistic and sensory traits, see Table 1. Future research with larger cohorts of autistic individuals (across a wider age range) would be beneficial for determining how perspective-taking and embodiment interacts with the development of language acquisition, theory of mind and sensory sensitivities in autistic individuals. Finally, we note that based on the current findings we cannot disambiguate between a mental rotation and a visuospatial transposition strategy. Future research should aim to understand in more detail what strategy is preferably engaged by autistic participants.

## Conclusion

Using MEG, we show that the neural basis of visual perspective-taking differs between autistic adolescents and non-autistic age-matched controls. Behaviourally, autistic participants took longer to respond, specifically when the angle between self and other perspectives was high. Neural processing in the autistic group was dominated by alpha-related signatures in the visual system. In contrast, the control group showed strong neural responses within an extended theta-band network of temporo-parietal and pre-frontal regions required for embodied transformation. We argue this represents a split strategy between the groups, with autistic adolescents appearing to favour a visuospatial transposition or mental object rotation strategy, whereas non-autistic adolescents seem to favour an embodied self-rotation strategy, similar to non-autistic adults (Kessler & Rutherford, 2010; Seymour et al., 2018; Wang et al., 2016). Our findings also support claims that ASD is solely associated with differences in perspective-taking (VPT-2) rather than tracking (VPT-1) (Pearson et al., 2013, 2016).

## Author Contributions

RS, KK & GR co-designed the study. RS collected the data, carried out the analysis, organised the data and code for sharing and was the primary author for this article. GGW helped with data collection. HW gave data analysis guidance.

## Data Availability Statement

The data that support the findings of this study are available on reasonable request from corresponding author, R.S., in a preprocessed and de-anonymized form. The raw data are not publicly available due to ethical restrictions. MATLAB data analysis code for this study will be made available openly on Github upon acceptance of the manuscript.

## Acknowledgements

We wish to thank the volunteers and their parents who gave their time to participate in this study.

## Supplementary Materials

**Supplementary Figure 1.**
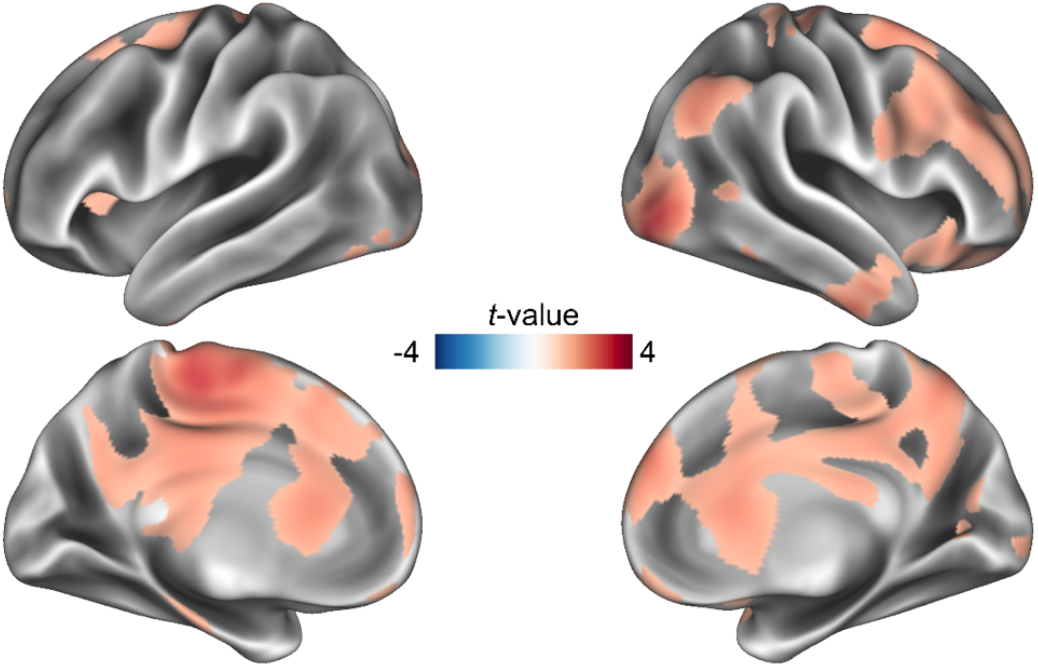
Perspective-taking theta source localisation results. For the non-autistic control group, we localised theta power (3-7 Hz) and statistically compared LR-160 versus LR-60 trials (0-0.65 s). Power maps are presented on an inflated brain from HCP Workbench. Only clusters passing a p<.05 threshold, corrected for multiple comparisons, are shown. In replication of Wang et al (2016) and Seymour et al (2018), significant clusters are observed in right posterior temporo-parietal junction, right visual ventral stream, right prefrontal cortex, and in medial cortical areas such as the cingulate cortex (ranging from posterior to anterior.

**Supplementary Figure 2.**
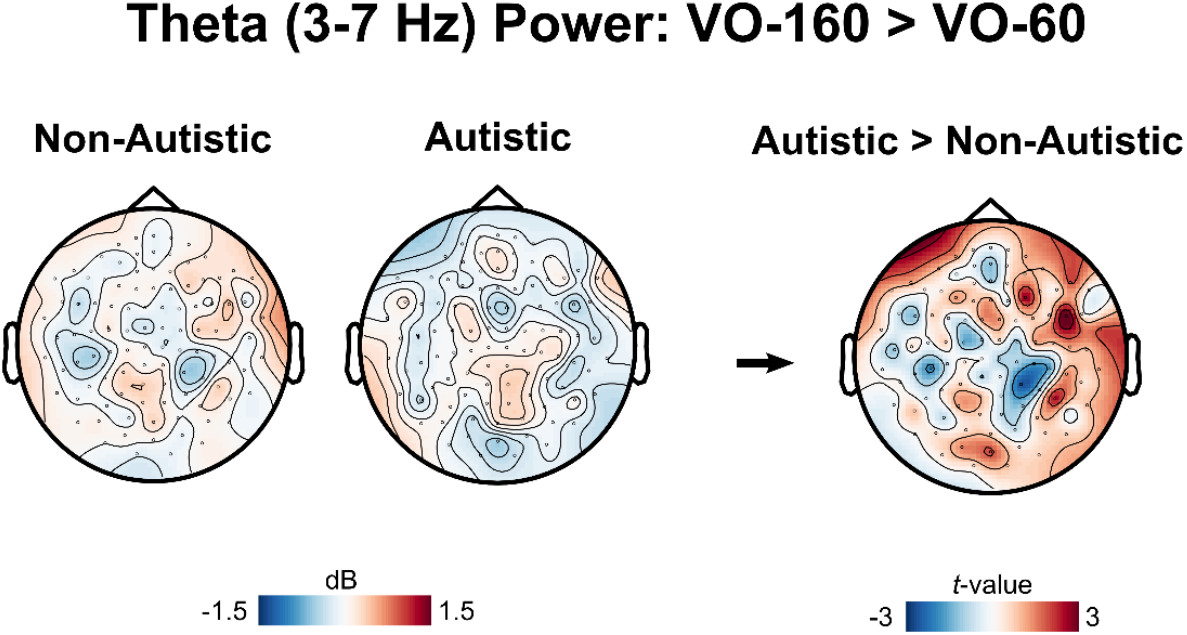
We investigated theta-power during perspective-tracking where participants were asked to judge if the target was visible or occluded from the avatar’s perspective. Sensor-level time-frequency representations were calculated using the same pipeline as outlined in the main manuscript. Paralleling the perspective-taking analysis, we compared theta power (3-7 Hz, 0-0.65s) in VO-160 vs. VO-60 trials. Overall there were very small changes in theta power and no significant group differences for perspective-tracking.

**Supplementary Figure 3.**
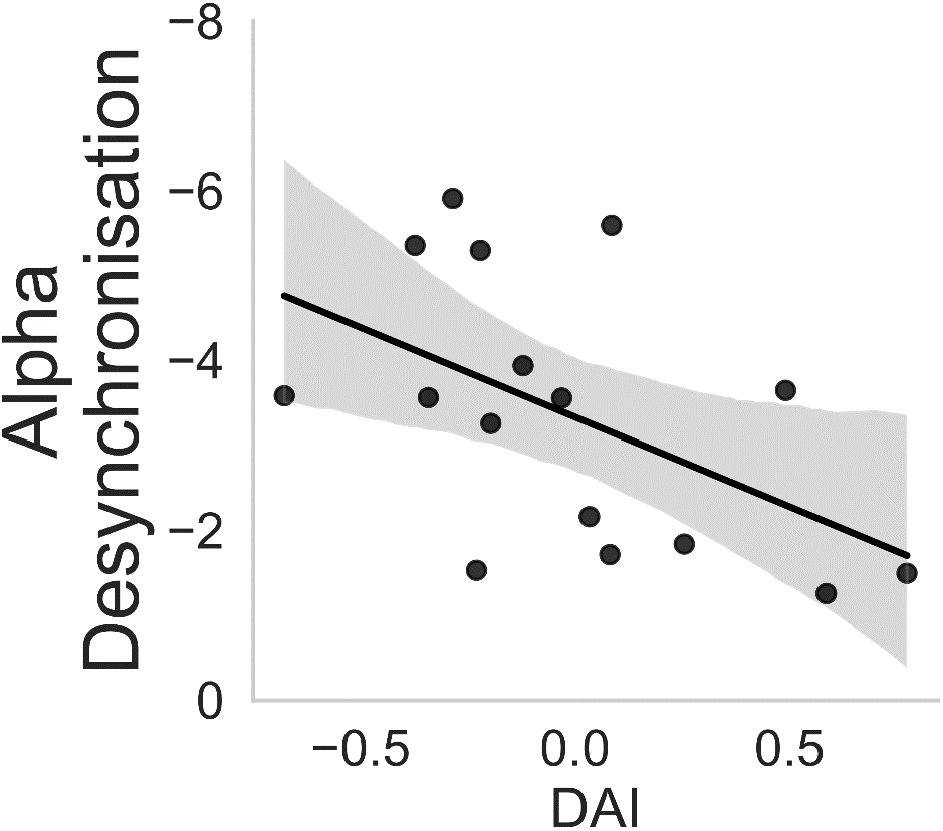
In the autistic group a significant correlation, r = .512, p = .043, was found between the amount of alpha-band synchronisation during perspective-taking and the directed asymmetry index (DAI) reported in Seymour et al., (2019) which represents a measure of alpha-band V4-to-V1 feedback connectivity in the visual system. In this instance, the lower the DAI value the greater the amount of feedback connectivity in the visual system, which correlates with the amount of alpha desynchronisation observed in the perspective taking task.

